# Right fronto-parietal networks mediate the neurocognitive benefits of enriched environments

**DOI:** 10.1101/2021.09.01.458571

**Authors:** Méadhbh B. Brosnan, Nir Shalev, Jivesh Ramduny, Stamatios N. Sotiropoulos, Magdalena Chechlacz

**Affiliations:** Department of Experimental Psychology, University of Oxford, Oxford, UK; Oxford Centre for Human Brain Activity, University of Oxford, Oxford, UK; Wellcome Centre for Integrative Neuroimaging, University of Oxford, Oxford, UK; Turner Institute for Brain and Mental Health, Monash University, Melbourne, Victoria, Australia; Sir Peter Mansfield Imaging Centre, School of Medicine, University of Nottingham, Nottingham, UK; National Institute for Health Research (NIHR), Nottingham Biomedical Research Centre, Queen’s Medical Centre, Nottingham, UK; Centre for Human Brain Health, University of Birmingham, Birmingham, UK; School of Psychology, University of Birmingham, Birmingham, UK

**Keywords:** cognitive ageing, reserve, alertness, superior longitudinal fasciculus, diffusion MRI

## Abstract

Exposure to enriched environments (EE) throughout a lifetime, providing so called reserve, protects against cognitive decline in later years. It has been hypothesised that high levels of alertness necessitated by EE might strengthen the right fronto-parietal networks (FPN) to facilitate this neurocognitive resilience. We have previously shown that EE offset age-related deficits in selective attention by preserving grey matter within right fronto-parietal regions. Here, using neurite orientation dispersion and density imaging (NODDI), we examined the relationship between EE, microstructural properties of fronto-parietal white matter association pathways (three branches of the superior longitudinal fasciculus, SLF), structural brain health (atrophy), and attention (alertness, orienting and executive control) in a group of older adults. We show that EE is associated with a lower orientation dispersion index (ODI) within the right SLF1 which in turn mediates the relationship between EE and alertness, as well as grey- and white-matter atrophy. This suggests that EE may induce white matter plasticity (and prevent age-related dispersion of axons) within the right FPN to facilitate the preservation of neurocognitive health in later years.

## INTRODUCTION

In the foreword to the 2020 world health organisation (WHO) landmark report on healthy ageing Dr Tedros Adhanom Ghebreyesus (WHO Director-General) states that “Humans now live longer than at any time in history. But adding more years to life can be a mixed blessing if it is not accompanied by adding more life to years.” An integral component of living life to the fullest in our later years is the capacity to maintain adequate cognitive abilities in older age. But human ageing is characterised by a sheer diversity of trajectories, with some older adults maintaining almost youth-like physical and cognitive capacities, and others experiencing frailty, disability, and dementia. Multiple modifiable factors across a lifetime have been shown to influence the trajectories of cognitive ageing (Livingston et al., 2020). Older adults who have engaged in cognitively and socially enriched environments exhibit greater resilience to cognitive decline when faced with neuropathological conditions like Alzheimer’s Disease (e.g., Stern et al. 1992; Bennett et al., 2006; Stern et al. 2018; Xu et al. 2019), a phenomenon referred to as neurocognitive reserve, cognitive reserve or simply reserve (see Cabeza et al., 2018; Stern et al, 2019; Cabeza et al., 2019; Stern et al., 2020). The concept of reserve originates from long-standing observations with Alzheimer’s patients whereby higher levels of education attainment delay the onset time of clinical symptoms of the disease and, consequently, cognitive decline (Stern et al., 1992; Stern, 2012; Stern et al., 2020). However, it is increasingly evident that such benefits to neurocognitive health are not exclusively obtained through education, but are additionally noted for leisure and social activities, and occupational engagements throughout a lifetime (e.g. Valenzuela & Sachdev, 2006; Opdebeeck et al., 2015). Correspondingly, it has been proposed that enriched, cognitively engaging environments across a lifetime contribute to optimal brain health and to prevent or at least delay the onset of dementia later in life (for review see Nelson et al., 2021; Cabeza et al., 2018). Yet despite compelling, large-scale longitudinal evidence for this phenomenon, we are just beginning to understand the neurobiological basis by which enriched environments impact the brain to cultivate resilience.

Enriched environments (EE) necessitate the continued engagement of several core cognitive processes, including alertness, sustained attention, and awareness, all of which rely on the right fronto-parietal networks (e.g., Beck et al., 2001; Naghavi and Nyberg, 2005; Hester et al., 2005; Singh-Curry and Husain, 2009; Stuss, 2011). Continued activation of these right lateralised networks has been theorised to cultivate and support the neuroprotective phenomenon of cognitive reserve (Robertson, 2014). The locus coeruleus norepinephrine (LC-NE) alertness system has strong projections to the right fronto-parietal networks (Oke et al. 1978; Robinson 1979; Grefkes et al. 2010; Jodo and Aston-Jones 1997; Jodo et al. 1998; Singewald and Philippu 1998; Shalev et al. 2019), and the strengthening of these networks by enriched environments over a lifetime is proposed to arise from the continued engagement of the LC-NE system over extended periods of time (obertson, 2014).

Numerous studies provide evidence that cognitive experiences (e.g., training or learning new skills) result in experience-dependent brain changes in brain structure in both the grey and white matter regions (i.e., structural plasticity changes; e.g., Draganski et al., 2004; Scholz et al., 2009; Woollett & Maguire, 2011). As such, the beneficial influence of a life-long exposure to EE on cognitive performance later in life may be understood in terms of structural plasticity processes which may optimize brain functioning and cognitive performance (Lövdén et al., 2010; Barulli & Stern, 2013) to offset neuroanatomical deficits relating to poor brain health (e.g., structural atrophy; Chan et al., 2018).

Our previous work using mathematical models of visual attention (Brosnan et al., 2017), causal manipulation techniques (Brosnan et al., 2017; Brosnan et al., 2018) and voxel based morphometry (Shalev et al., 2020) provide increasing support for the proposal that EE may facilitate neuroprotective resilience through specifically impacting the structural (grey matter volume) and function (lateralised asymmetry of visual processing speed) of right hemisphere fronto-parietal regions (see also Van Loenhoud et al 2017; and see Robertson 2013, 2014 for a detailed review theorising right lateralised underpinnings of reserve). Cortical regions within the fronto-parietal networks are connected by white matter association pathways comprising three branches of the superior longitudinal fasciculus (SLF1, SLF2 and SLF3; Thiebaut de Schotten et al., 2011). In younger adults, inter-individual differences in the ability to efficiently attend and respond to sensory information are mediated by variability in white matter organisation of the superior longitudinal fasciculus (Chechlacz et al., 2015a; Marshall et al., 2015; Brosnan et al., 2020). Our recent work suggests that EE offsets age-related deficits in selective attention by preserving grey matter within right fronto-parietal regions. Here we test whether white matter microstructure of the right SLF might be cultivated by EE to facilitate better maintenance of attention function in older adults.

White matter pathways underpin the efficiency of communication between discrete cortical regions. Physical characteristics, including micro- and macrostructural properties of the white matter (including volume, axonal diameter, myelination, fiber coherence and neurite density) influence both the neurophysiological function of the tract and functional connectivity, with subsequent consequences for behaviour (e.g., Chechlacz et al., 2015b, Marshall et al., 2015; Deligianni et al., 2016; Brosnan et al., 2020). White matter changes associated with the ageing process affect connectivity within neural networks and underlie gradual age-related cognitive decline (e.g., Raz & Rodrigue, 2006; Barrick et al., 2010; Burzynska et al., 2010; Westlye et al., 2010). In contrast to invasive animal studies and post-mortem anatomical and histological human studies, microstructural properties of white matter and age-related white matter changes can be non-invasively studied in the living human brain with diffusion magnetic resonance imaging (diffusion MRI; for recent review see Lerch et al., 2017). While diffusion MRI only provides indirect estimates of biological properties of the white matter, new developments in acquisition and data modelling increasingly enable more robust and biologically plausible measures of white matter microstructural properties (Assaf & Basser, 2005; Zhang et al., 2012; Lerch et al., 2017; Alexander et al., 2019). Diffusion tensor imaging (DTI) based on conventional single-shell acquisition protocols and DTI-derived microstructural measures of white matter properties, such as fractional anisotropy (FA) and mean diffusivity (MD), have been widely employed to study age-related white matter changes (e.g., Sexton et al., 2014; Barrick et al., 2010; Westlye et al., 2010; Burzynska et al., 2010). However, while DTI-derived measures are indeed sensitive to age-related alterations in white matter microstructure, these are nonspecific measures, which cannot be attributed to specific changes in tissue microstructure (Pierpaoli et al., 1996; Beaulieu, 2009). For example, age-related reduction in FA might be a result of decreased neurite density, changes in fibre orientation dispersion and/or other changes such as in the degree of myelination (Beaulieu, 2009, Zhang et al., 2012). By contrast multi-shell acquisition protocols combined with biophysically plausible models have been shown to provide more specific estimates of the microstructural properties of white matter tissue (Lerch et al., 2017). One such approach is Neurite Orientation Dispersion and Density Imaging (NODDI; Zhang et al., 2012), which has been previously used to provide detailed accounts of white matter changes associated with development, ageing and several neurological disorders (e.g., Kodiweera et al., 2016; Kunz et al., 2014; Billiet et al., 2015; Chang et al., 2015; Adluru et al., 2014; Billiet et al., 2014; Winston et al., 2014; Timmers et al., 2015; Veldsman et al., 2020). NODDI-derived parameters, intra-cellular volume fraction (ICVF) and the orientation dispersion index (ODI), respectively measure neurite packing density and dispersion of neurites/axons (an estimate of fiber coherence). These two parameters are effectively independent features encoded in the FA, both providing a more biologically specific model of observed changes, which could not previously be disentangled from FA measures derived from DTI. For example, previous research suggests that the increase in FA during development is driven by increasing neurite density, while the reduction in FA later in life is driven by the increase in ODI (Chang et al., 2015). Moreover, changes in NODDI-derived parameters have been shown to be more sensitive to ageing and predictive of cognitive performance (Merluzz et al., 2016; Kodiweera et al., 2016).

In the current study we use neurite orientation dispersion and density imaging (NODDI) to examine the relationship between microstructural properties of the SLF, enriched environments (measured by proxies such as education, professional, leisure and social activities), attentional capacity (representing cognitive processes contributing to reserve), and structural brain health (grey and white matter atrophy) in a group of older adults. To assess attention function, we used a well-established task (the Attention Network Test; ANT; Fan et al., 2005; Petersen & Posner, 2012). The ANT has been developed to measure three partially distinct ‘networks’ supporting attention: *Alerting* attention in response to a temporally predictive cue, *Orienting* attention in space, and exerting *Executive Control* to resolve conflict and enhance relevant information. To corroborate the proposal that cognitively stimulating environments and experiences across the lifetime strengthen the right lateralised fronto-parietal networks to facilitate neurocognitive health later in life, we hypothesise that EE induces white matter plasticity (prevents age-related dispersion of neurites and/or reduction in neurite density) within the right SLF to facilitate better maintenance of attention function (specifically alertness) and superior brain health (less volume atrophy) in older adults.

## METHODS

### Participants

A total of 50 older adults (22 males; age range 65-84; mean ± SD age 73.5 ± 4.7) were recruited to the study, which consisted of behavioural testing (with the Cognitive Reserve Index questionnaire and the ANT task) and a magnetic resonance imaging (MRI) session. All participants were recruited either from the Psychology panel of elderly volunteers, or the Birmingham 1000 Elders group, both established at the University of Birmingham. The two panels of elderly volunteers consisted of adults aged 65 or over who were in good health and had no pre-existing cognitive impairment. All study volunteers had normal or corrected-to-normal vision, had no history of psychiatric or neurological disease and identified as right-handed. The study was approved by the University of Birmingham Ethical Review Committee. All study participants provided written informed consent and received monetary compensation for participation in agreement with approved ethics protocol.

### Environmental Enrichment

To estimate levels of lifelong exposure to enriched environments, all study participants completed the Cognitive Reserve Index questionnaire (CRIq; Nucci et al. 2012). This is a semi-structured interview, assessing educational attainment, along with the complexity of professional engagements and a wide variety of leisure and social activities. The CRIq measure comprises of 3 subscales (CRI education, CRI working activity and CRI leisure time), and a composite score (overall CRIq; Nucci et al. 2012). Each of the three subscales are derived based on both the frequency and duration (in weeks, months, or years) of the various activities across lifespan.

### Attentional Capacity

#### The Attention Network Test

We used the Attention Network Test (ANT; Fan et al., 2002), which has been employed extensively to study attention among older adults (e.g., Fernandez-Duque and Black, 2006; Jennings et al., 2007; Ishigami and Klein, 2011; McDonough et al., 2019). The task was designed to test the efficiency of the three ‘networks’ supporting attention: *Alerting* (achieving and maintaining a state of alertness), *Orienting* (the selection of information from sensory input), and *Executive control* (resolving conflict among competing response dimensions; Posner and Petersen, 1990). In the task, outlined in Figure 1, partisans are requested to judge the direction of a central arrow (target) surrounded by four flanking stimuli (two on each side). The target and its surrounding flakers appear in one of two locations; above or below a fixation mark at the centre of the monitor along the central vertical line. The flankers can be either neutral flankers (lines with no arrow heads), congruent (arrows pointing in the same direction as the target), or incongruent (arrows pointing in the opposite direction to the central target). Each trial starts with a variable fixation period (400-1600ms; FP) and the presentation of the flanked target is preceded by one of four cueing conditions (see Figure 1). These are: a Central Cue, where a central warning signal (an asterisk) is presented; a Double Cue, where two asterisks are presented indicating the two possible locations of the flanked target along the horizontal line; a Spatial Cue indicating where the target will appear using a single asterisk; and No Cue. The cue is always presented for 100ms and followed by another fixation period of 400 ms. A fixation cross remains at the centre of the screen throughout the trial, with the exception of the Central cue condition where it is replaced for 100 ms by an asterisk. The target surrounded by flankers is always presented either until the participant responds (RT) or, in the case of no response for a maximum of 1700 ms. In total each trial lasts for 4000 ms (Figure 1). Thus, once the target and flankers disappear, the next trial is preceded by a variable fixation interval based on the duration of first fixation (FP) and reaction time (RT).

**Figure 1.**
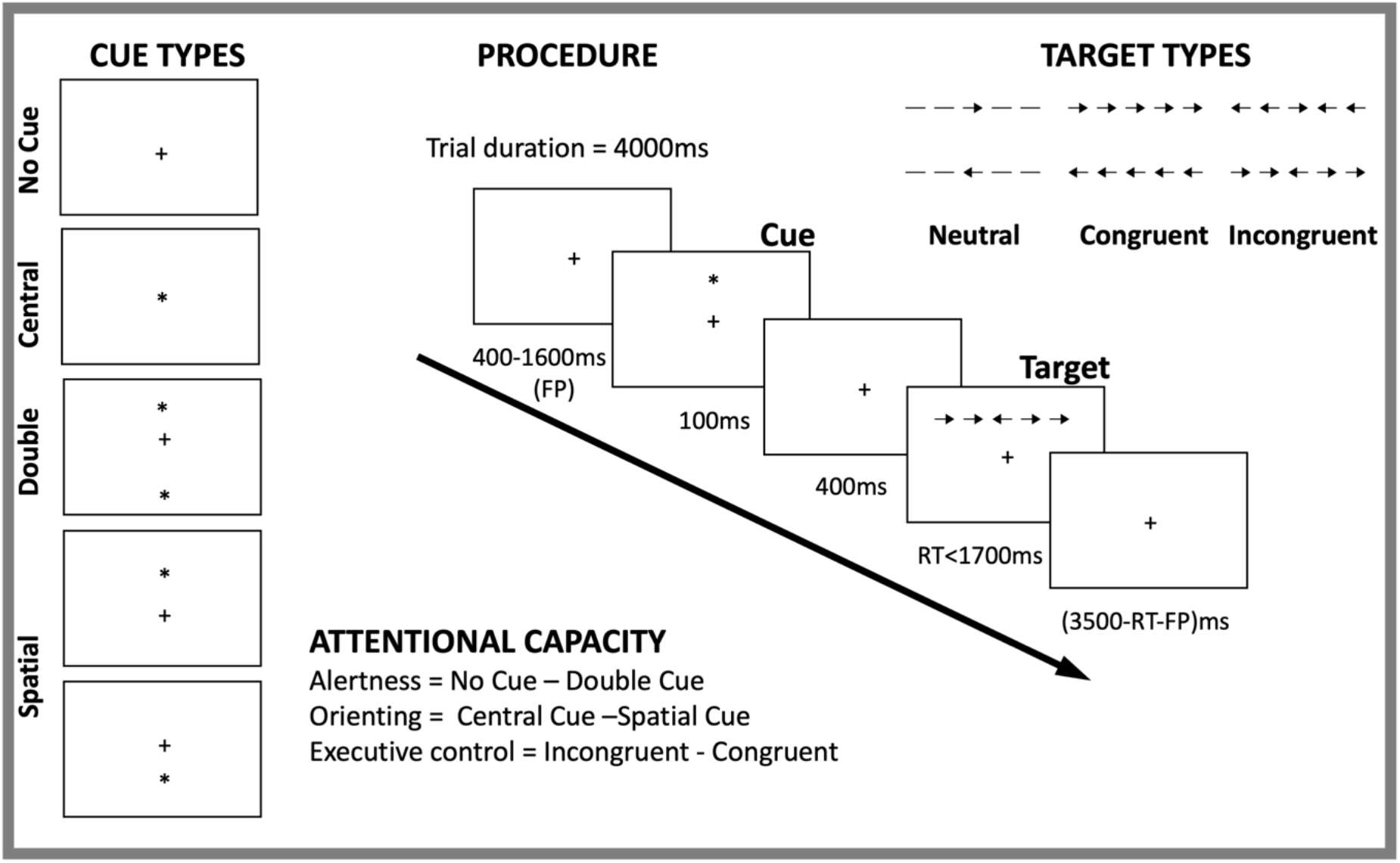
Illustration of the attention network test (ANT) procedure, including types of target and cue as well as example of a trial and estimation of attention capacity parameters.

#### ANT Task Procedure

Participants were instructed to attend to the central fixation cross and to use the mouse buttons to indicate the direction of the flanked arrow when presented (i.e., left mouse click for leftward arrowheads, and right mouse click for rightward arrowheads). The various conditions (four different cues and three levels of congruency) were randomised within blocks. The task began with 24 practice trials with feedback, followed by 3 blocks without feedback. Within each block there were 96 trials, presented in a random order. These consisted of Cue Condition (x4), Target Locations (x2), Target Directions (x2), Flanker Conditions (x3), Repetitions (x2). The practise block took approximately 2 min and each subsequent experimental block took approximately 5min to complete.

#### Measuring Attentional Capacity (ANT Scores)

We derived the ANT scores (alertness, orienting and executive control/conflict) as per previously published work (e.g., Fan et al., 2002; Posner, 2008) by calculating the difference between mean reaction time (RT) of the different conditions (defined by either cue or target type in the ANT procedure). The alertness score was calculated by subtracting mean RT in the double-cue condition (i.e., the two warning cues corresponding to the two possible target locations) from the mean RT in the no-cue condition (*Alerting = RT_no cue_ – RT_double cue_*). As such, the Alerting score indicates the degree to which an individual uses the external cues to benefit their reaction times. Larger alerting scores indicate a relative difficulty in maintaining alertness in the absence of an external cue i.e., decreased ability to rely on internal (or intrinsic) alertness (Posner, 2008). However, it is possible that larger alertness scores might instead indicate a more efficient use of the cue. One way to dissociate the two is by testing for a correlation between the *Alerting* score and the mean reaction time from the no cue condition. A positive association would imply that participants who relied more on the alerting cue to facilitate performance were slower, thereby indicative of decreased intrinsic alertness capacities. The orienting is calculated by subtracting the mean RT in the spatial-cue condition from the mean RT in the central-cue condition (*Orienting = RT_central cue_ – RT_spatial cue_*). The spatial cue, which is always valid, provides information about subsequent target location and thus facilitate orienting attention before target arrival. As such, the Orienting Score reflects the difference between responses to targets that follow spatially predictive and non-predictive cues. Larger Orienting scores indicates a better capacity to orient attention and select sensory input (Fan et al., 2002). Finally, the executive control score is calculated by subtracting mean RT in the congruent condition from the mean RT in incongruent condition across all cue types. (*Executive control/Conflict = RT_incongruent_ – RT_congruent_*). The Conflict score represents the capacity to resolve response conflict to targets that appear among distractors. The larger Conflict score indicates less ability (difficulty) in resolving conflict (Posner, 2008). We present descriptive statistics and estimate the Pearson correlations among the four task indices: Alerting, Orienting, Conflict and mean reaction time (RT; Fig. 3A, C).

### Fronto-parietal Microstructure

#### MRI data acquisition

T1-weighted scans (MPRAGE with spatial resolution 1×1×1mm^3^) and multi-shell diffusion-weighted images were acquired at the Birmingham University Imaging Centre (BUIC) using a Philips 3T Achieva MRI system with a 32-channel head coil. The multi-shell diffusion acquisition comprised a single-shot EPI, 2×2×2mm^3^, 5 × b=0 s/mm^2^, 50 × b=1000s/mm^2^, 50 × b=2000s/mm^2^ and, in order to correct for susceptibility-induced artifacts 5 × b=0 s/mm^2^ (Andersson et al., 2003). T1-weighted scans were acquired with the following parameters: 176 slices, TR = 7.5 ms, TE = 3.5 ms and flip angle = 8°. Diffusion-weighted scans were acquired with the following parameters: 56 slices, TR=9000ms, TE=81.5 and flip angle = 90°.

#### T1-weighted data pre-processing and total grey matter volume estimation

T1-weighted scans were pre-processed using the UK Biobank T1-weighted pipeline (Alfaro-Almagro et al., 2018), which was employed to correct for bias fields, apply skull-stripping and align data to the MNI152 standard space, before segmenting the T1 images into different tissue classes, i.e., grey matter (GM), white matter (WM) and cerebrospinal fluid (CSF) using FAST (FMRIB’s Automated Segmentation Tool Zhang et al. (2001). These data were subsequently used to calculate total GM and total WM volume in mm^3^, normalized for head size using SIENAX package (Smith et al., 2002).

#### Diffusion data pre-processing, microstructural model fitting and SFL tractography

Diffusion-weighted scans were pre-processed using the UK biobank pipeline (Alfaro-Almagro et al., 2018; Figure 2). The T1-weithed pre-processing was employed to correct for bias fields, apply skull-stripping and align data to the MNI152 standard space. The diffusion-weighted pre-processing was applied to correct for susceptibility induced distortion, eddy-current distortion and participant movement induced distortions using the EDDY toolbox (Andersson & Sotiropoulos, 2016) and to obtain transformations of the diffusion to structural and standard space. Subsequently, we applied the NODDI (Neurite Orientation Dispersion and Density Imaging) model to the multi-shell EDDY-corrected diffusion data (Zhang et al., 2012, Tariq et al., 2016) using the cuDIMOT (CUDA Diffusion Modelling Toolbox; https://users.fmrib.ox.ac.uk/~moisesf/cudimot/; Hernandez-Fernandez et al., 2019) to estimate voxel-wise microstructural parameters, including intra-cellular volume fraction (ICVF), and orientation dispersion index (ODI). In addition, a diffusion tensor model (Basser et al., 1994) was fitted to low b-value (b=1000s/mm^2^) shells of the EDDY-corrected diffusion data to obtain fractional anisotropy (FA) maps for each participant. Next, we performed automated probabilistic tractography using predefined protocols for identifying major WM tracts in the left and right hemispheres, as described in FSL’s XTRACT tool (https://fsl.fmrib.ox.ac.uk/fsl/fslwiki/XTRACT; de Groot et al., 2013; Warrington et al., 2020). Prior to running XTRACT, we fitted the crossing fibre model (FSL’s BEDPOSTX; Behrens et al., 2007) to each subject’s data to estimate up to three fibre orientations per voxel, and ran nonlinear transformations to the MNI152 standard space. XTRACT automatically reconstructs a set of predefined white matter pathways, including the three branches of the superior longitudinal fasciculus (SFL 1,2,3). These three tracts were reconstructed and used for the purpose of data analysis. The SLFs’ tract probability density maps, normalized by the total number of valid streamlines, were thresholded at 0.1% and binarised to produce a tract mask for each tract in standard space. Finally, for each participant we calculated set of image derived phenotypes (IDPs), characterizing different microstructural properties of the three branches of SLF. Specifically, for each tract (left and right SLF1,2,3), and for each DTI (FA) and NODDI (ODI, ICVF) parameter, the weighted mean value (i.e., the mean weighted by the tract probability in each voxel) of the parameters within given tract was calculated. Outliers were assessed defined for the neuroimaging measures as individuals more than 3 times the IQR and three outliers were removed.

**Figure 2.**
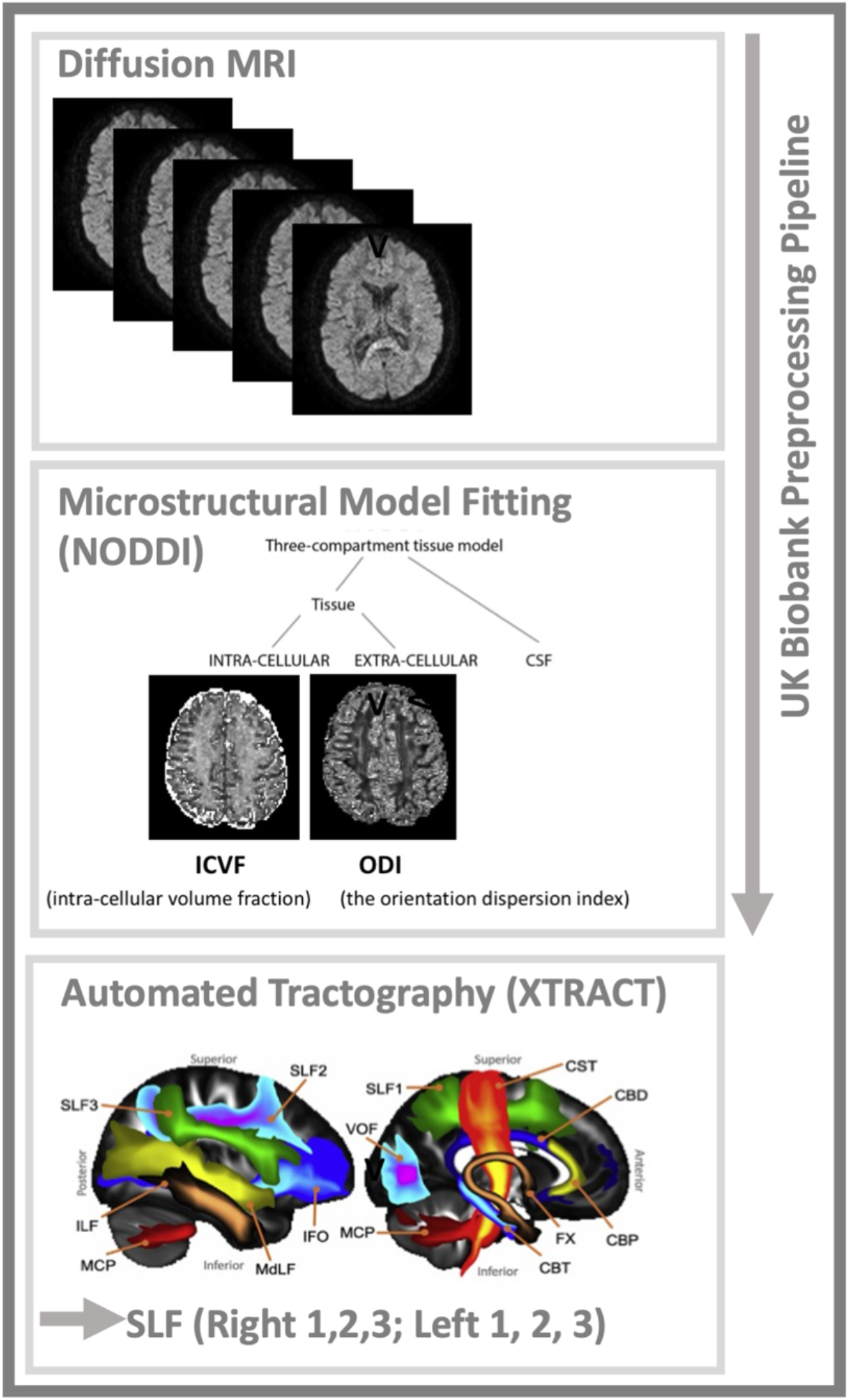
Overview of the diffusion MRI data analysis pipeline.

### Statistical Analyses

To assess whether environmental enrichment was differentially associated with hemisphere-specific microstructural organisation of the SLF we modelled EE (the composite CRIq score) as a function of three separate microstructural estimates: orientation dispersion index (ODI), intra-cellular volume fraction (ICVF), and fractional anisotropy (FA) within the three SLF branches in each hemisphere. For each model, age was entered as a nuisance covariate in the first step, then the ODI estimates of the six SLF branches (i.e., branches 1-3 for both hemispheres) were entered in the model using a stepwise approach. These analyses revealed that ODI (within the right SLF1) was particularly sensitive to the impact of EE. In the analyses above EE was estimated using the composite CRIq score. In order, to identify whether specific types of enrichment were driving the observed effect of EE on the right SLF1, a follow up analysis of the three subscales (Education, Occupational, and Leisure engagements) was performed.

To investigate whether ODI within the SLF was associated with the degree of grey- and white-matter volume atrophy (GMVa and WMVa respectively) in the older individuals’ brains, we modelled total GM and total WM volume (normalized for head size), as a function of ODI within the three SLF branches in each hemisphere. Again, age was entered as a nuisance regressor in step 1 of both models, and the ODI estimates for all six SLF branches were entered into each model with a stepwise approach. These analyses identified a specific association between both GMVa, and WMVa with ODI within the right SLF1. We subsequently tested whether a causal association existed between EE, ODI within the right SLF1, and brain atrophy (separately GMVa and WMVa) using bootstrapped mediation analyses. For this, bootstrapped mediation analyses (5000 samples) were performed using the PROCESS computational toolbox (Hayes, 2012; 2013). More specifically, the plausibility of a causal model was investigated whereby EE (predictor variable X) causally influenced white matter microstructure (ODI) of the right SLF 1 (mediator variable M), to in turn exert a causal influence over GMVa and WMVa in the older adults (outcome variables Y). The confidence intervals (Cis) reported for the indirect effect are bootstrapped Cis, based on 5000 samples and are considered significant when they do not contain zero.

Finally, to determine whether the right SLF1 showed a meaningful relationship to attentional capacity, we next modelled ODI within this tract as a function of the three ANT scores (alertness, orienting, executive control) using a stepwise regression model, again with age as a nuisance covariate in the first step of the model. As we observed an association between ODI within the right SLF1 and alertness, we subsequently tested whether a causal association existed between EE, ODI within the right SLF1, and alertness using bootstrapped mediation analysis as described above (causal mediation model EE-> rSLF1 -> Alertness). To explore which aspect of EE was driving our effects, a follow up mediation analysis with the three subscales (Education, Occupational, and Leisure engagements) of CRIq was performed.

## RESULTS

### Attentional capacity: performance on the attention network test (ANT)

Overall mean accuracy was high (mean accuracy 98%; SD=.02%; range 92% - 100%) indicating that participants did not have any difficulty completing the task. Our analysis focused on RT-based indices measuring individual differences in attentional capacity i.e., three network scores used to represent the efficiency of the alerting, orienting and executive control (Fan et al, 2002; Posner, 2008). Figure 3A summarizes RT data averaged for each target and cueing condition.

**Figure 3.**
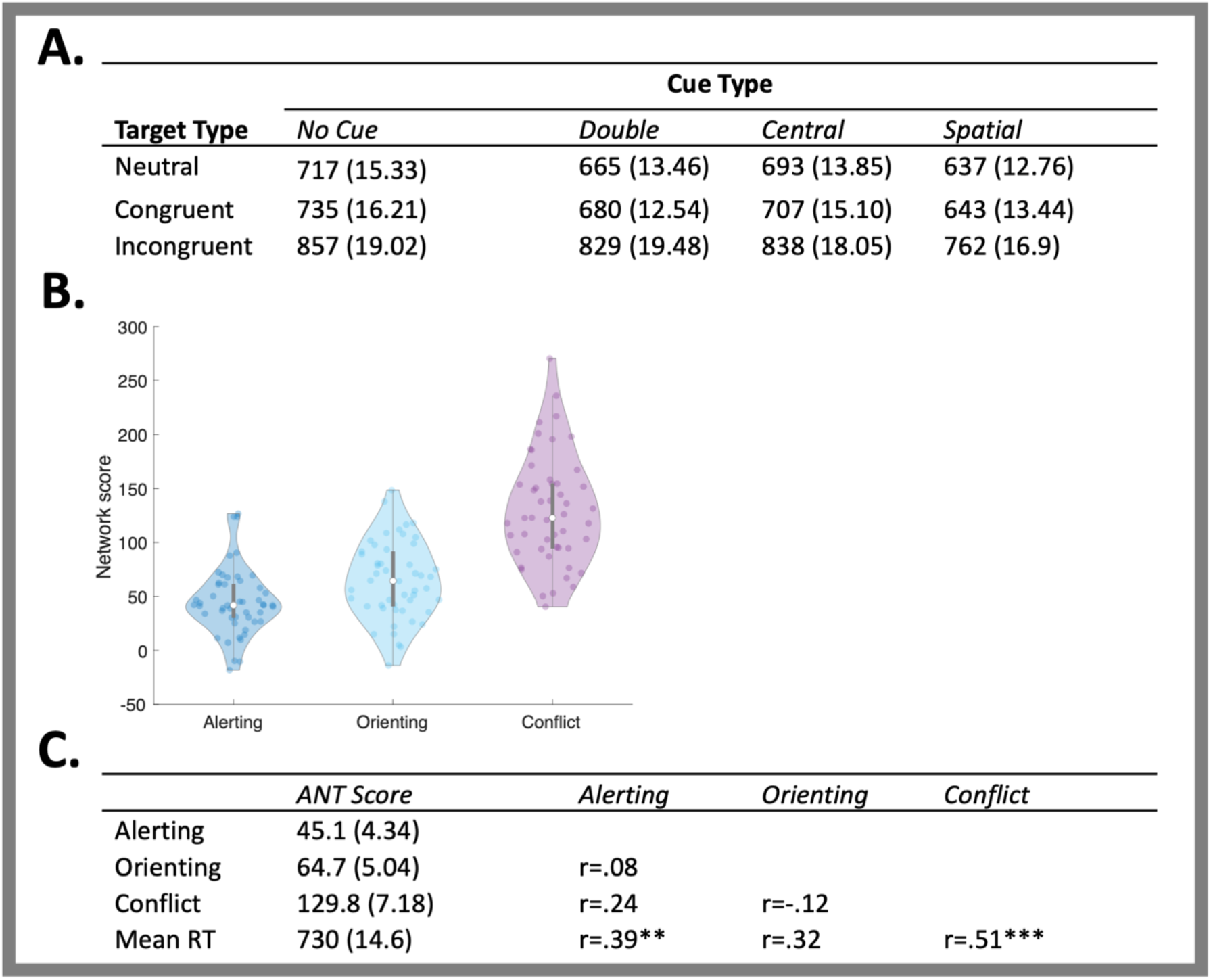
Performance on the ANT (raw scores and the derived *network* scores; Alerting, Orienting, and Conflict). **A**. The mean RT (standard error) for each target and cueing condition. **B.** The Alerting, Orienting, and Conflict network scores based on the ANT performance are plotted to illustrate attentional capacity distribution in the studied group of *N*=50 older adults. **C.** The mean network scores (effect), mean RT (standard error) and correlations between attention networks. *Note* *** denotes *p*<.0005, ** denotes *p*<.005

The distribution of the network scores is depicted in Figure 3B. The scattered data points illustrate that the majority of the observations were positive, thus indicating that in this cohort of older adults, most participants (a) benefitted from the alerting cue with larger scores arising from decreased ability to rely on alertness; (b) had good capacity to orient attention i.e., benefitted from a spatial cue (orienting attention) prior to the arrival of target; and (3) were better at judging congruent targets, compared to incongruent i.e., had decreased ability (difficulty) in resolving conflict.

As suggested in the Methods section, the interpretation of the ANT alertness measure is not straightforward as a larger alerting score could be indicative of either decreased ability to rely on internal alertness or more efficient use of a cue. We tested the correlation between the Alertness score and the mean RT in the “No cue” condition and found a significant high correlation (r=0.56, p<0.0005. Accordingly, people who benefitted more from the alerting cue were also relatively slow when there was no cue. This is in line with the former interpretation of the Alertness index such that in our sample higher scores were associated with lower level of internal alertness.

The mean network scores (effect), mean RT and correlations between these indices are shown in Fig. 3C. The correlation analyses were conducted to examine independence of the attentional capacity measures. As in originally published ANT paper (Fan et al., 2002) reporting no significant correlations between network scores, we found no correlations between alerting, orienting and executive control scores indicating the independence of estimated measures of alertness, orienting and executive control. Mean RT correlated with the alerting and conflict scores. Note, three participants were excluded in the MR analysis as outliers. All subsequently are conducted with a sample of N=47.

### EE mitigates neurite (axonal) dispersion in the right SLF1

We modelled EE (the composite CRIq score) as a function of orientation dispersion index (ODI) within the SLF branches (SLF1,2 and 3) in each hemisphere (Figure 4). Age was entered as the first step in the model and offered no significant improvement in model fit over the intercept only model (*R*^2^_change_ =02, *F*_change_=.94, *p F*_change_ =.34, unstandardized beta =.67, SE=.68, t=.97, *p*=.34). Next, ODI estimates for each of the six SLF branches were entered into the model using a stepwise approach. This model offered a significant improvement in fit (*R*^2^_change_ =.12, *F*_change_=5.86, *p F*_change_ =.02) and led to the inclusion of the right SLF1 (t=−2.42, *p*=.02, Fig. 4), and exclusion of all other SLF branches (all *t*>−1.50, p>.14), indicating that greater EE was associated with less dispersion of neurites (lower ODI) within this tract. To obtain accurate parameter estimates of this model, not influenced by other uninformative variables, we modelled EE directly as a function of the right SLF1 and report the results in Table 3. We next assessed whether EE affects neurite density (as measured by intra-cellular volume fraction, ICVF) of the SLF using a similar modelling approach, this time with ICVF within the SLF branches as predictor variables. We found no evidence that EE was associated with altered neurite density (ICVF) in any of the SLF branches (all *t*<1.10, *p*>.28), suggesting that EE specifically altered neurite dispersion and not neurite density within the right SLF1. Finally, using the same modelling approach we assessed whether EE varied as a function of fractional anisotropy (FA). We found no evidence that EE was associated with altered FA within any of the SLF branches (all *t*<.33, *p*>.13).

**Figure 4.**
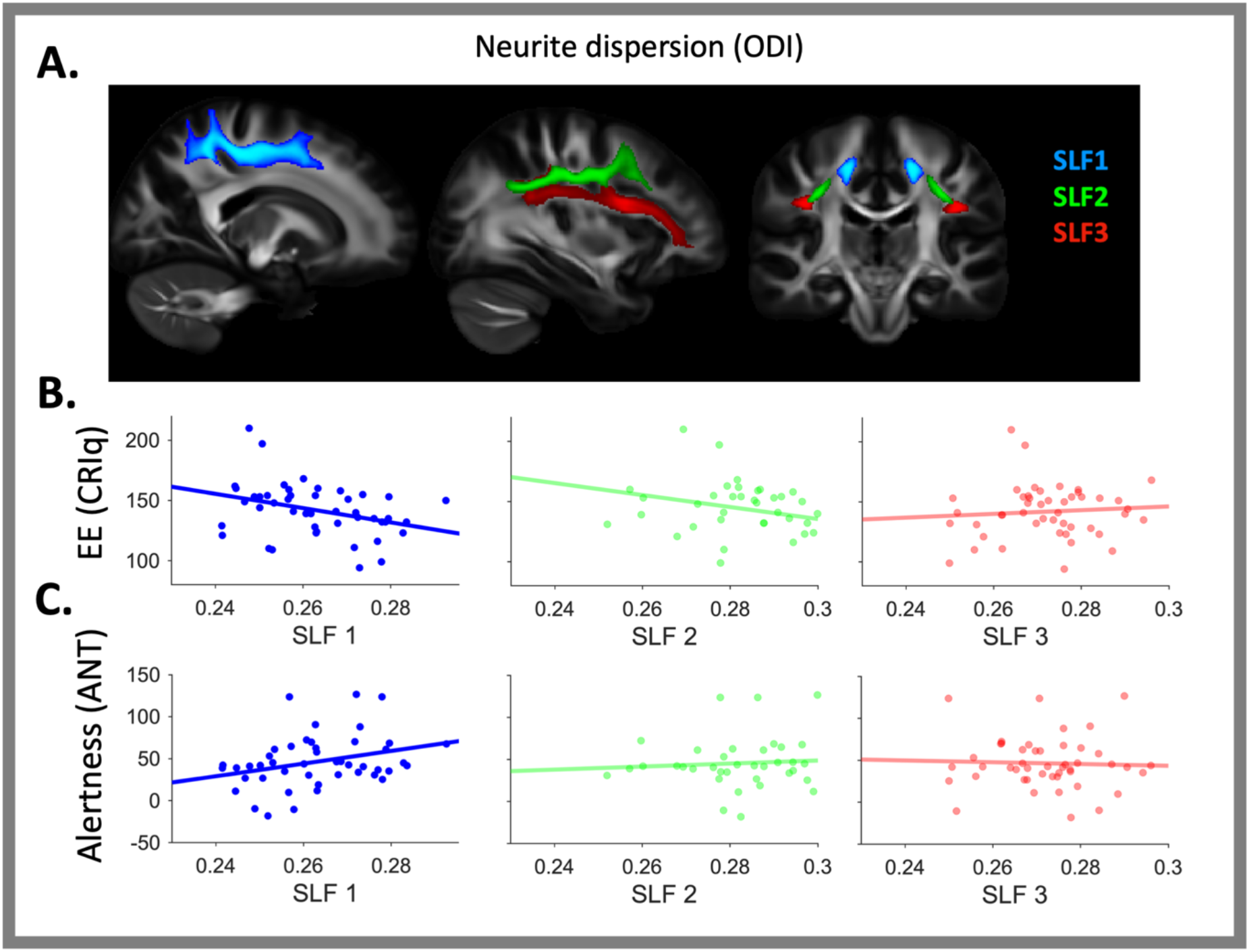
**(A)** XTRACT reconstructed SLF branches. The relationship between neurite dispersion (ODI) within the right SLF and EE (**B**) and Alertness (**C**).

**Table 3.**
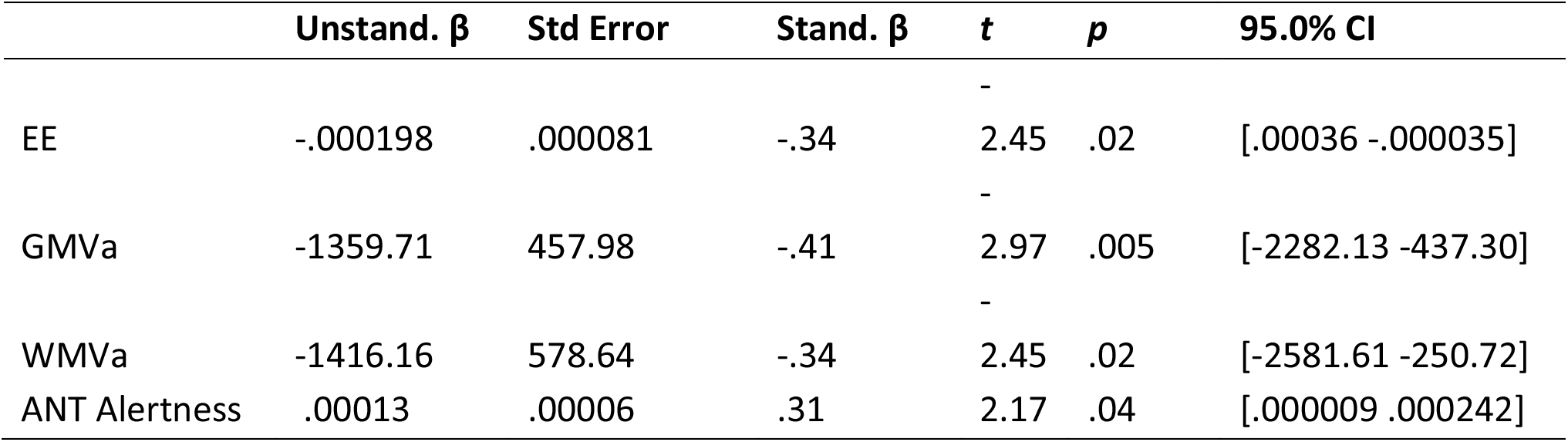
Neurite Dispersion of the right SLF 1 as a function of EE, Grey Matter Volume Atrophy (GMVa), White Matter Volume Atrophy (WMVa), and Alertness

### The effect of EE on the right SLF1 is driven by professional engagement

Having observed that EE was associated with altered neurite/axonal dispersion within the right SLF1, we next asked the question of whether this effect was driven by any specific facet of enriching activities. Our assessment of EE (the CRIq) comprised of three subscales; leisure, work, and education. As such, we explored the direct association between the right SLF1 and all three subscales using a model with right SLF1 as the dependent variance, age as a nuisance predictor in the first stage of the model, and the three CRIq subscales as predictors in the second stepwise stage of the model. This revealed a specific association between the work subscale of the CRI and the right SLF1 such that greater occupational complexity and professional engagements were associated with decreased dispersion of neurites (ODI) in the older adults (*t*=−3.50, *p*=.001). No such associations were observed for either leisure activities or education (both *t*<.05, *p*>.95).

### The right SLF1 mediates the association between EE and Structural Brain Health (grey- and white-matter atrophy)

To test whether ODI of the SLF relates to structural health of the ageing brain, both grey-matter and white volume (both normalised by head size) were modelled separately as a function of age and the six SLF tracts (SLF 1-3, left and right hemisphere). For grey matter atrophy (GMVa) when age was entered in the first step of the model, it offered no significant improvement in model fit over the intercept only model (R^2^_change_ =.01, F_change_=.47,p F_change_ =.5, unstandardized beta =−.91, SE=1.33, t=−.68, p=.50). Next, ODI estimates for each of the six SLF branches were entered into the model of grey matter volume using a stepwise approach. This model offered a significant improvement in fit (R2_change_ =.17, F_change_=8.85, p F_change_ =.005) and led to the inclusion of the right SLF1 (t=−2.98, p=.005), and exclusion of all other SLF branches (all t<.55, p>.58), indicating that less dispersion of neurites (i.e., lower ODI) within the right SLF1 was associated with less grey matter atrophy (i.e., greater total grey matter volume). To obtain accurate parameter estimates of this model, not influenced by other uninformative variables, we modelled GMVa directly as a function of the right SLF1 and report the results in Table 3.

For white matter atrophy (WMVa), again age was entered as the first step in the model and offered no significant improvement in model fit (R^2^_change_ =.01, F_change_=.34, p F_change_ =.56, unstandardized beta =−.95, SE=1.64, t=−.58, p=.56). Adding the ODI estimates for the six SLF branches using a stepwise approached offered a significant improvement in fit (R^2^_change_ =.12, F_change_=5.99, p F_change_ =.02) and again led to the inclusion of the right SLF1 (t=−2.45, p=.02), and exclusion of all other SLF branches (all t<.45, p>.41), indicating that less dispersion of neurites (lower ODI) of the right SLF1 was associated with less white matter atrophy (i.e., larger total white matter volume). To obtain accurate parameter estimates of this model, not influenced by other uninformative variables, we modelled WMVa directly as a function of the right SLF1 and report the results in Table 3.

Our results so far indicate that EE is associated with less neurite dispersion (ODI) of the right SLF1, and that less ODI within this specific tract is associated with less atrophy in both grey and white matter. This raises the possibility that a causal relationship may exist such that the extent to which EE positively impacts structural brain health (i.e., offsets white and grey matter atrophy) is dependent, at least partly, on the degree to which EE has altered white matter properties of the right SLF1. To formally test this hypothesis, we ran two causal mediation model (EE → ODI rSLF1 → GMVa; EE →ODI rSLF1 → WMVa). Bootstrapped mediation analyses showed an indirect effect for both GMVa (Direct effect = .31, p=.28; Indirect effect .23, 95% CI [.04 .48]) and WMVa (Direct effect =.24, p.51; Indirect effect .25, 95% CI [.01 .54]). This indicates that the degree to which EE positively mitigates grey and white matter atrophy of the ageing brain is dependent in part, on the degree to which EE has altered neurite dispersion properties of the right SLF1 (Figure 5).

**Figure 5.**
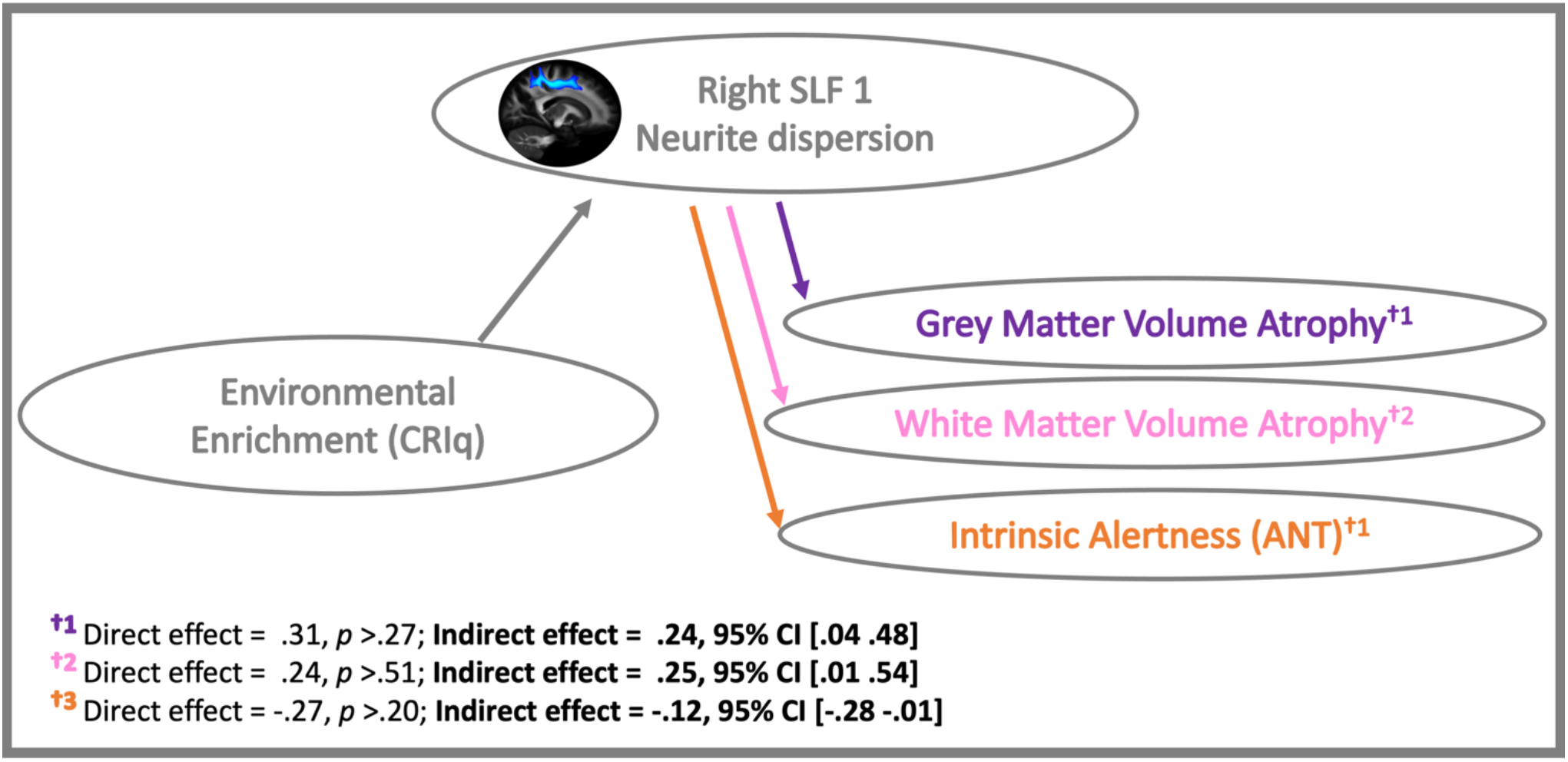
Causal mediation model showing an indirect relationship between EE and Alertness in older adults, which is mediated by neurite dispersion within the right SLF1.

### Neurite dispersion in the right SLF1 is associated with alertness score

The results thus far demonstrate that EE facilitates superior maintenance of neurite (axonal) dispersion in older adults within the right SLF1. This suggests that plasticity of the right SLF1 might be specifically sensitive to the beneficial effects of environmental enrichment on ageing brain health. To assess whether the right SLF1 showed a meaningful relationship to alertness, here we modelled this tract as a function of the three subscales of the ANT; Alertness, Orienting, Conflict, see Fig. 4C). Age was entered as the first step in the model and offered no improvement on the intercept only model (*R*^2^_change_ =0, *F*_change_=.02, *p F*_change_ =.88, unstandardized beta =0, SE=0, t=−.16, *p*=.88). Next, we modelled the right SLF1 as a function of the three ANT subscales using a stepwise approach. This led to a significant improvement in model fit (*R*^2^_change_ =.09, *F*_change_=4.59, *p F*_change_ =.04) with neurite dispersion of the right SLF1 uniquely associated with the Alertness (*t*=2.14, *p*=.04) but not Orienting (t=−1.00, *p*=−32) or Conflict (*t*=−1.63, *p*=.11) components of the task. To obtain accurate parameter estimates for the relationship between neurite dispersion of the right SLF1 and ANT alertness, we modelled the direct association without other uninformative signals, and report the results in Table 3.

### The right SLF1 mediates the association between EE and alertness

The findings thus far suggest that a lifetime of EE is associated with reduced neurite (axonal) dispersion within the right SLF1 in older adults. In addition, we observed that reduced right SLF1 neurite dispersion is consequential to cognition, such that it facilitates better maintenance of internal alertness in advanced age (Figure 4B). This raises the possibility of a causal association between these factors such that EE facilitates alertness by impacting plasticity of the right SLF1. To test this hypothesis, we ran a causal mediation model; EE→ ODI rSLF1 −→Alertness (Figure 5). Bootstrapped mediation analyses (5000 samples) revealed a significant mediation such that EE was associated with superior white matter microstructure (reduced neurite dispersion) within the right SLF1 which, in turn, facilitated better internal alertness. We note there was no direct relationship between EE and alertness suggesting that the association between these two variables is dependent on white matter properties of the right SLF1 (Direct effect = −.27, p=.21; Indirect effect −.1172, 95% CI [−.2837 −.0098]). In accordance with the findings above which highlighted a privileged association between occupational engagements and ODI of the right SLF1, repeating the mediation model with the work subscale (as opposed to the composite CRIq score) additionally resulted in a significant mediation effect (Direct effect = −.1924, p=.44; Indirect effect −.1777, 95% CI [−.4018 −.0185]).

## DISCUSSION

In the current study, we examined how microstructural properties of the SLF varied according to exposure to enriched environments (EE) to subsequently influence markers of neurocognitive health in older adults. Our findings indicate that greater exposure to enriched environments across a lifetime is associated with reduced axonal dispersion within the right SLF1 in older adults. This in turn mitigates declines in structural brain health (assessed via grey- and white-matter volume atrophy) and intrinsic alertness (captured with the well-validated ANT test). To the best of our knowledge, this is the first study providing direct evidence linking microstructural properties of a long-range association pathway within the right fronto-parietal networks (the right SLF1) to behavioural markers of neurocognitive reserve and brain health. Here we discuss how these results corroborate our previous work (Brosnan et al., 2017; Brosnan et al., 2018, Shalev et al., 2020) and that of others (Van Loenhoud et al. 2017), to provide further experimental evidence in support of the theory that enriched environments strengthen right hemisphere fronto-parietal networks to facilitate the phenomenon of cognitive reserve (Robertson 2013, 2014).

The findings presented here indicate that microstructure of the right SLF1 might be specifically sensitive to the beneficial effects of environmental enrichment to neurocognitive health in older adults. Brain matter volume, indicative of age-related atrophy, is often used as a proxy of ageing brain health (Stern, 2012; Lövdén et al., 2010; Barulli & Stern, 2013; Cabeza et al, 2018; Chan et al., 2018). Previous theoretical as well experimental work suggests that lifelong exposure to cognitive and social enrichment, providing so called cognitive reserve, can promote healthy cognitive function despite objective markers of vulnerability in the brain including atrophy (e.g., Chan et al., 2018) or disease-related neuropathological changes (e.g., Alzheimer’s; Xu et al., 2018). Yet the mechanisms by which this reserve is facilitated have remained unclear (Cabeza et al., 2018; Stern et al., 2019; Husain, 2021). Here we observed that the extent to which neurite (axonal) dispersion is preserved within the right SLF1 is directly associated with both structural brain health (grey- and white-matter atrophy) and cognitive function (intrinsic alertness on the ANT) in later years. More specifically, causal mediation models indicated that the associations between EE with both structural brain health and alertness were dependent on neurite dispersion of the right SLF1. This suggests that the benefits of EE to neurocognitive health in older adults depend, at least in part on the degree to which white matter microstructural architecture within the right SLF1 has been altered.

Novel and complex environments require high levels of alertness, which in turn optimise the processing upcoming signals (Petersen & Posner, 2012). Functionally linked with noradrenaline (Aston Jones & Cohen 2005; Marrocco & Davidson 1998), alertness has been associated with right-lateralised fronto-parietal networks and thalamic structures (Posner & Petersen, 1990; Corbetta & Shulman, 2002; 2011; Strum & Willmes 2001; Singh-Curry & Husain, 2009; Fan et al 2005; see also Coull et al., 2000; Fan et al 2005), with deficits in maintaining alertness reported following right but not left hemispheric strokes (Posner et al., 1987; Posner, 2008). Given the critical relevance of noradrenaline to neurocognitive resilience (Robertson, 2013, 2014) alertness is an ideal candidate to facilitate the beneficial consequences of EE. At the neural level, it is likely that repeated exposure to EE necessitates high levels alertness which strengthens right hemisphere fronto-parietal networks, to exert beneficial effects on cognitive symptoms of ageing. Our findings suggest a self-perpetuating relationship between enriched environments (assessed by CRIq) which likely necessitate high levels of alertness, well-established structural and functional networks which support the cognitive capacity of alertness (right SLF1), and intrinsic alertness capacities (assessed via the Alertness subscale of the ANT). This is not only directly in line with the hypothesis that alertness, through its association with the right lateralised noradrenergic system and the right fronto-parietal networks, is a critical determinant of cognitive reserve (Robertson, 2014) but also support the rationale for designing interventions to maximally modulate alertness in older adults.

The superior longitudinal fasciculus (SLF) is a white matter fronto-parietal association pathway which consists of three branches (Thiebaut de Schotten et al., 2011). The most dorsal branch (the SLF1) has projections to the frontal eye fields (FEF), premotor cortex, and the intralparietal sulcus (IPS; Stuss et al., 2002). These regions comprise a functional network often referred to as the dorsal attention or dorsal fronto-parietal network (Fox et al., 2006; Corbetta & Shulman, 2002; Vincent et al., 2008; Corbetta et al. 2008). This network is activated by internal goals and expectations and links the processing of sensory information, including the formation of perceptual decisions, with the relevant motor commands (Corbetta et al., 2008; Brosnan et al., 2020). As such, it is likely that white matter architecture within the dorsal SLF facilitates the coordination and communication of information between sensory, decision, and motor regions to subsequently influence the efficiency of an individual’s response (Brosnan et al., 2020). Our findings may suggest that a lifetime of intrinsic alertness ‘training’, afforded by high EE, may induce white matter plasticity within the right SLF1 to facilitate better intrinsic alertness in older years. In our cohort, these effects were driven by the complexity of professional and occupational engagements. Future work in larger cohort studies should investigate what are the optimal factors for inducing white matter plasticity within the right SLF1 and whether, in vulnerable cohorts, novel intervention techniques (e.g., brain stimulation and neurofeedback) could be used to induce similar changes to structural organisation within this pathway. Addressing these hypotheses would provide increasing support for the utility of the right SLF as a marker of resilience in older adults, and facilitate translational avenues to optimise interventions aimed at preventing age-related cognitive decline.

The results presented in this manuscript raise that intriguing possibility that white matter plasticity within the right fronto-parietal networks may be induced by enriched environments to facilitate resilience to cognitive decline later in life. Here we show that EE was associated with decreased dispersion of axons (lower ODI) specifically within the right SLF1, but not the other SLF branches. Previous studies contrasting younger and older participants have indicated that increased dispersion of axons is one of the indicators of age-related brain changes indicative of poorer cognitive function (Kodiweera et al., 2016; Nazeri et al., 2015; Billiet et al., 2015; Chang et al., 2015). Moreover, previous reports assessing the effect of age on white matter have demonstrated that NODDI-derived measures outperform measures derived from DTI (e.g., Kodiweera et al., 2016). In agreement with Kodiweera et al., our data similarly point to greater sensitivity of ODI compared to DTI-derived FA, such that our associations between the SLF and behaviour were observed specifically for ODI and not FA. Our findings are in line with growing evidence from both rodent and human studies that experience and training induces white matter plasticity, reflected through changes in white matter microstructural properties in adult brain (e.g. Scholz et al., 2009; Blumenfeld-Katzir et al., 2011, Sampaio-Baptista et al., 2013). One of the proposed mechanisms underlying this white matter plasticity are activity-dependent changes in myelination (for review see Sampaio-Baptista and Johansen-Berg, 2017). Myelination and the NODDI-derived parameters used here are somewhat related but predominantly complementary measures of white matter microstructure (Fukutomi et al, 2019; Billiet et al., 2015). Thus future investigations with both animal and human participants should disentangle the precise biological basis underpinning the impact of a lifetime of EE on neurocognitive health. It should be also noted here that despite improved estimations of microstructural features of the white matter with NODDI compared to DTI, ODI (axonal dispersion) measures, which estimates the degree of fiber coherence, could be still affected by crossing with other white matter tracts. As such, while our results point to an association between EE and plasticity of the right SLF1, it is a possibility that our findings could be driven, at least to some extent by changes to other right fronto-parietal pathways which cross the SLF 1.

One caveat to our findings is that factors which may have drawn individuals to high levels of EE throughout their lifetime (including socioeconomic demographics, personality, genetic, and early life cognitive abilities) were not available for investigation. As such we cannot rule out the possibility that alternative mediating factors are contributing to our observed associations between environment, the brain, and behaviour. However, several points suggest this was not the case. First, previous work with monozygotic (identical) twin studies has shown EE positively impacts cognition over and above genetic factors (Lee et al. 2013). Moreover, our effects of EE were driven by the professional and not education subscale of the CRIq, suggesting that early life cognitive engagement was not the critical influence in this cohort. Finally, increasing evidence suggests that the ageing brain maintains sufficient plasticity such that it can be targeted in later years to increase reserve and resilience (Peeters et al. 2020; Lenehan et al. 2016; Edwards et al. 2017).

Although the beneficial effects of EE for neurocognitive resilience are well established, the mechanisms by which this occurs are still under considerable debate. The findings presented here, in concert with our previous findings (Brosnan et al. 2017; Shalev et al. 2020) suggest that anatomical properties within the right fronto-parietal networks might be neural correlates of resilience in the ageing brain. In further support of this hypothesis, brain stimulation targeting the right fronto-parietal networks not only improves behavioural and EEG markers of attention but also temporarily alters the lateralised impact of a lifetime experiences in low reserve individuals such that it resembles that of their high reserve peers (Brosnan, Arvaneh, et al. 2018; Brosnan, Demaria, et al. 2018). Although neuropathological markers of Alzheimer’s Disease are associated with cognitive deficits, over 50% of the variability in symptomatology still remains unexplained (Boyle et al., 2018; Boyle et al. 2021). We propose that white matter properties of the right FPN be tested as objective markers of resilient neurocognitive ageing that might help capture the heterogeneity associated with the clinical presentation of Alzheimer’s disease and be used to assay the efficacy of interventions aimed at ameliorating age-related cognitive decline.

## ACKNOWLEDGMENTS

We warmly thank the volunteers from the School of Psychology panel and the Birmingham 1000 Elders group for participation in this study. This work was supported by the Birmingham-Nottingham Strategic Collaboration Fund (BNSCF336 to SNS and MC) and by a Wellcome Trust Institutional Strategic Support Fund critical data award (204846/Z/16/Z to MC). MC was supported with a BRIDGE (Birmingham-Illinois Partnership for Discovery, Engagement and Education) Fellowship and MBB was supported with a Marie Skłodowska-Curie Fellowship from the European Commission (AGEING PLASTICITY; grant number 844246). This work was additionally supported by the NIHR Oxford Health Biomedical Research Centre and the Wellcome Centre for Integrative Neuroimaging was supported by core funding from the Wellcome Trust (203139/Z/16/Z).

## Notes

**CONFLICT OF INTEREST:** The authors declare no potential conflict of interest.

### Competing Interest Statement

The authors have declared no competing interest.

